# Interplay of geometry and mechanics in epithelial wound healing

**DOI:** 10.1101/2024.04.08.588496

**Authors:** Nandhu Krishna Babu, M Sreepadmanabh, Sayantan Dutta, Tapomoy Bhattacharjee

## Abstract

Wound healing is a complex biological process critical for maintaining an organism’s structural integrity and tissue repair following an infection or injury. Recent studies have unveiled the mechanisms involving the coordination of biochemical and mechanical responses in the tissue in wound healing. In this article, we focus on the healing property of an epithelial tissue as a material while the effects of biological mechanisms such as cell crawling and tissue proliferation is minimal. We present a mathematical framework that predicts the fate of a wounded tissue based on the wound’s geometrical features and the tissue’s mechanical properties. Precisely, adapting the vertex model of tissue mechanics, we predict whether a wound of a specific size in an epithelial monolayer characterized by certain levels of acto-myosin contractility and cell-cell adhesion will heal (i.e., close), shrink in size, or rupture the tissue further. Moreover, we show how tissue-mediated mechanisms such as purse-string tension at the wound boundary facilitate wound healing. Finally, we validate the predictions of our model by designing an experimental setup that enables us to create wounds of specific sizes in MDCK monolayers. Altogether, this work sets up a basis for interpreting the interplay of mechanical and geometrical features of a tissue in the process of wound healing.

## INTRODUCTION

The natural healing process of wounded tissues is a multifaceted biological phenomenon crucial for preserving organismal structural integrity and facilitating tissue repair post-infection or injury. Research in this area has extensively focused on the repair of a damaged epithelial layer utilizing both *in vivo* platforms—such as sea urchin coelomocytes, fruit fly wing disks, and jellyfish medusae [1–5]—as well as *in vitro* model systems like monolayers of kidney epithelial cells (MDCK) [6–9], keratinocytes (HaCaT), and colon epithelial cell lines (CaCo-2) [10–15]. Biophysical factors such as cell-substrate interactions, mobilization of leading-edge cells, interaction of cells and extracellular matrix (ECM), and tissue’s response to growth factors dynamically interact to orchestrate different processes involved in wound healing [14, 16–18]. For instance, cells adjacent to the wound area exhibit purse-string like actin filament bundles which coupled with force-generating myosin motor proteins ultimately lead to wound closure. Concurrently, cells in the bulk tissue showcase coordinated migratory motion (crawling) through lamellipodial protrusions which also contribute towards the wound healing process [7, 8, 12, 19]. Notably, physical constraints such as the shape of the migrating front and wound geometry exert profound effects on wound repair [20, 21]. Despite the reasonable control provided by these experimental platforms over several biophysical cues, deciphering the relative contributions of different factors in the wound healing process has remained challenging.

On the other hand, over the last few decades, mathematical models have been developed to comprehensively describe epithelial tissue mechanics by elucidating the relative contribution of different factors involved in the wound healing process [22–27]. Some of these models treat epithelial tissue as a continuum thin layer, where individual cells are not distinguishable [12, 28–30]. While these models elucidate tissue-level dynamics, including cell sheet elasticity, tissue rheology, or tissue-substrate interaction, they fail to capture mechanics at a single-cell level, such as cell-cell adhesion and tissue intercalation. Alternatively, the vertex model represents epithelial tissue as a network of polygons (cells), with vertices moving subject to various microscopic forces arising from parameters such as cell elasticity, acto-myosin contractility, interfacial tension, and cell motility [1, 31–47]. This approach has been used extensively in describing the process of wound healing [1, 27, 48–52].

Although several studies have both computationally and experimentally investigated integrated mechanisms of wound healing such as cell crawling and tissue proliferation, the healing property of epithelial tissue as a material has been understudied. In this article, we focus on the role of different mechanical forces in the epithelial tissue such as cell-cell adhesion, acto-myosin contractility, and cell-wound interfacial tension in determining wound closure. We look at how the coordination and competition of these forces decide the fate of a given wound. Specifically, we present a model to calculate the energy of a wounded epithelial monolayer utilizing the 2D vertex model description of a tissue. Next, based on energy minimization our model predicts the fate of a wound—whether the wound closes (heals), shrinks, or ruptures the tissue. To study the effect of these forces on wound healing in isolation, we do not consider cell crawling, cell division, cell death, or neighbour exchange. We are specifically focusing on the final state of the wound and contribution of different mechanical forces in determining that state. We emphasize that the coarse-grained as well as static nature of this model allows us to do a parameter scan in the high dimensional space spanning a range of mechanical properties as well as geometric features of the tissue. Finally, we design an experimental setup using MDCK monolayers to closely replicate the physical scenario depicted in our model. We then validate our theoretical predictions qualitatively and quantitatively.

## MODEL DESCRIPTION

We begin our analysis by defining the energy of a single cell in the vertex model [37, 46], that describes the mechanics of an epithelial monolayer:

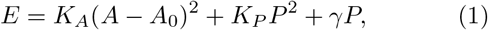

where, *K*_*A*_ represents the cell’s resistance to volume change and is related to the cell’s bulk elasticity modulus, *A* is the area of the cell, *A*_0_ is the preferred area of the cell, *K*_*p*_ represents the elastic constant of the contractile acto-myosin cortex, *P* is the perimeter of the cell and *γ* represents cell-cell interfacial tension. Next, we define a non-dimensional energy 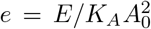, which can be expressed as

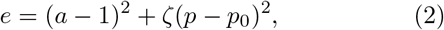

subject to an additive constant. Here, *a* = *A/A*_0_ is the area of a cell normalized to its preferred area, *ζ* = *K*_*P*_ */K*_*A*_*A*_0_ is a non-dimensional parameter set by the ratio of elasticity of ac<u>to-</u>myosin cortex to the bulk modulus of the cell, 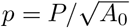 and 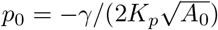, are the actual and preferred perimeters of the cell respectively, normalized to the characteristic length of the cell. The preferred perimeter of the cell *p*_0_ is set by the competition of the interfacial tension between the cells and the contractility of the acto-myosin cortex.

We consider a homogeneous epithelial monolayer consisting of *N* cells with an average area equal to the preferred area of the cell (i.e., ⟨*a*⟩ = 1). In this monolayer, a wound is modeled by removing a fraction *f*_0_ of the tissue, leaving *N* (1 − *f*_0_) cells in the tissue. If the instantaneous area fraction of the wound in this tissue is *f* , the average normalized area of each cell is ⟨*a*⟩ = (1− *f* )*/*(1 −*f*_0_). The average normalized perimeter of the cells is 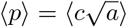 , where *c* is the shape factor. Along the perimeter of the wound, an effective non-dimensional cell-wound interfacial tension (purse-string tension) Γ replaces cell-cell interfacial tension 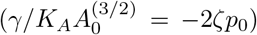. The normalized perimeter for a circular wound is 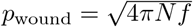. Under these considerations and neglecting the variance of cell-sizes, we can estimate the total energy of the tissue with an initial wound fraction *f*_0_ and instantaneous wound fraction *f* as,

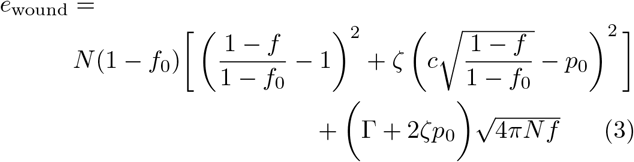

Eq. 3 allows us to predict the mechanical response of the wounded tissue for specific mechanical and interfacial properties (i.e., Γ, *ζ, p*_0_), tissue size (*N* ), and the initial size of the wound (*f*_0_). We evaluate the energy of the wounded tissue for different area fractions *f* (0 *< f <* 1) and decide its steady state by finding the area fraction *f*_ss_ corresponding to the minimum energy of the tissue [Fig. 1(b)-(d)]. Based on the value of the *f*_ss_, we can distinguish the mechanical response of the tissue into three categories: (i) if *f*_ss_ = 0, the wound completely closes [Fig. 1(b)]; (ii) if *f*_ss_ *< f*_0_, the wound shrinks but doesn’t close [Fig. 1(c)]; (iii) if *f*_ss_ *> f*_0_, the wound further expands (i.e., the tissue ruptures) [Fig. 1(d)].

**FIG. 1.**
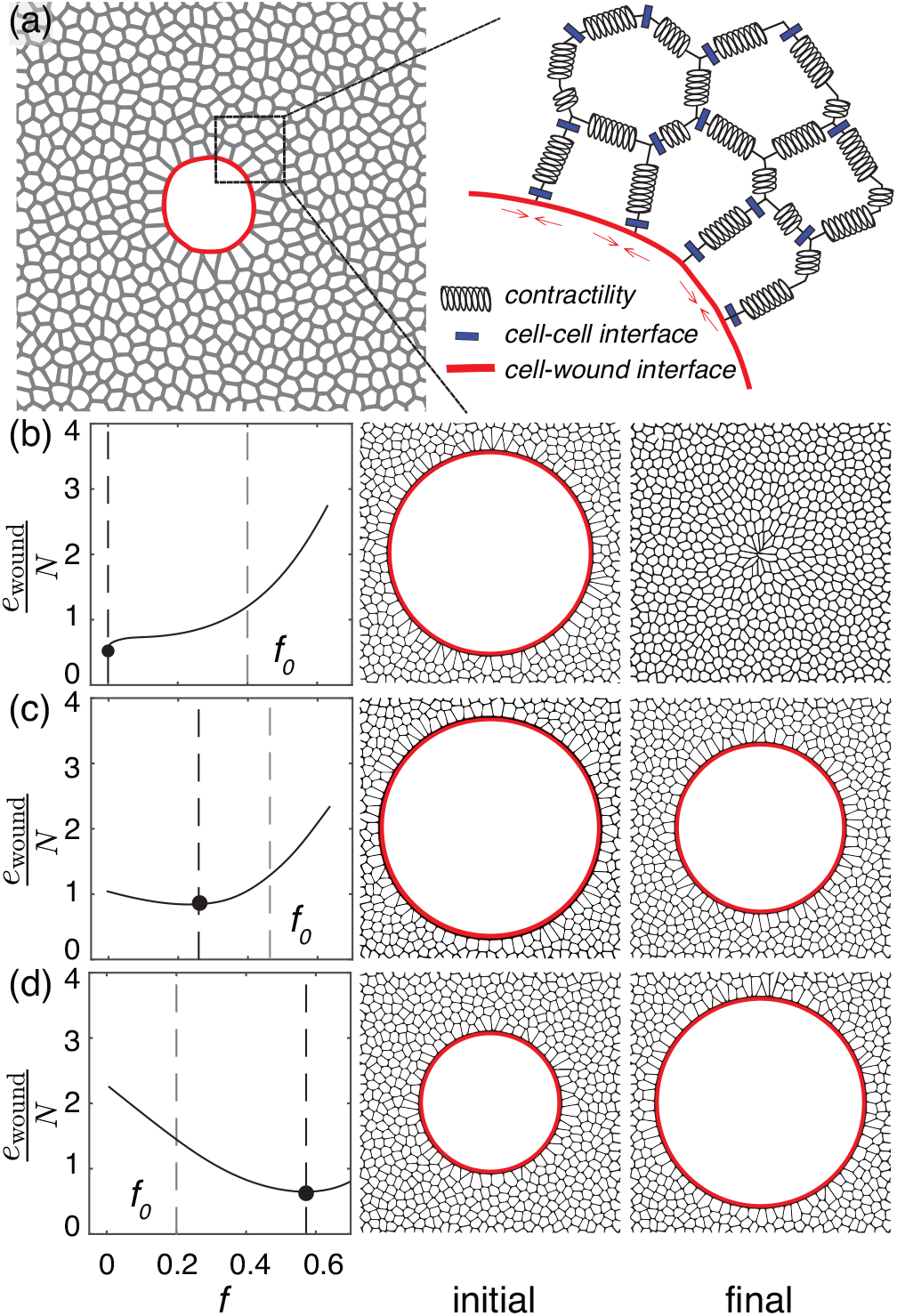
(a) Schematic of a wounded epithelial monolayer (left) along with a zoomed in version of the same schematic (right) that shows different relevant forces. (b)-(d) shows three fates of a wound: closure, shrinkage, and expansion respectively. (left) Normalized total energy of the tissue (*e*_wound_*/N* ) as a function of wound area fraction (*f* ) for three fates. The dashed grey and black lines represent the initial wound area fraction (*f*_0_) and the final wound area fraction (*f* ) corresponding to the minimum of *e*_wound_*/N* . (center and right) Schematic of the initial and final states of the wound.

## RESULTS

### Model Prediction

Three fates of the wound partition the space of geometric and mechanical parameters that describe the wounded monolayer (i.e., *p*_0_, *ζ*, Γ, *N* , *f*_0_). We vary the shape factor *c* between 3.67 (corresponds to a regular heptagon) and 3.81 (corresponds to a regular pentagon), describing the typical shape of the cells in an epithelial monolayer [37]. We mark the regions of the parameter space in grey as shown in Fig. 2(a) and Fig. 3, where the predicted mechanical response of the tissue is not uniform in this range of *c*. We keep *ζ* = 1 for the results presented here.

**FIG. 2.**
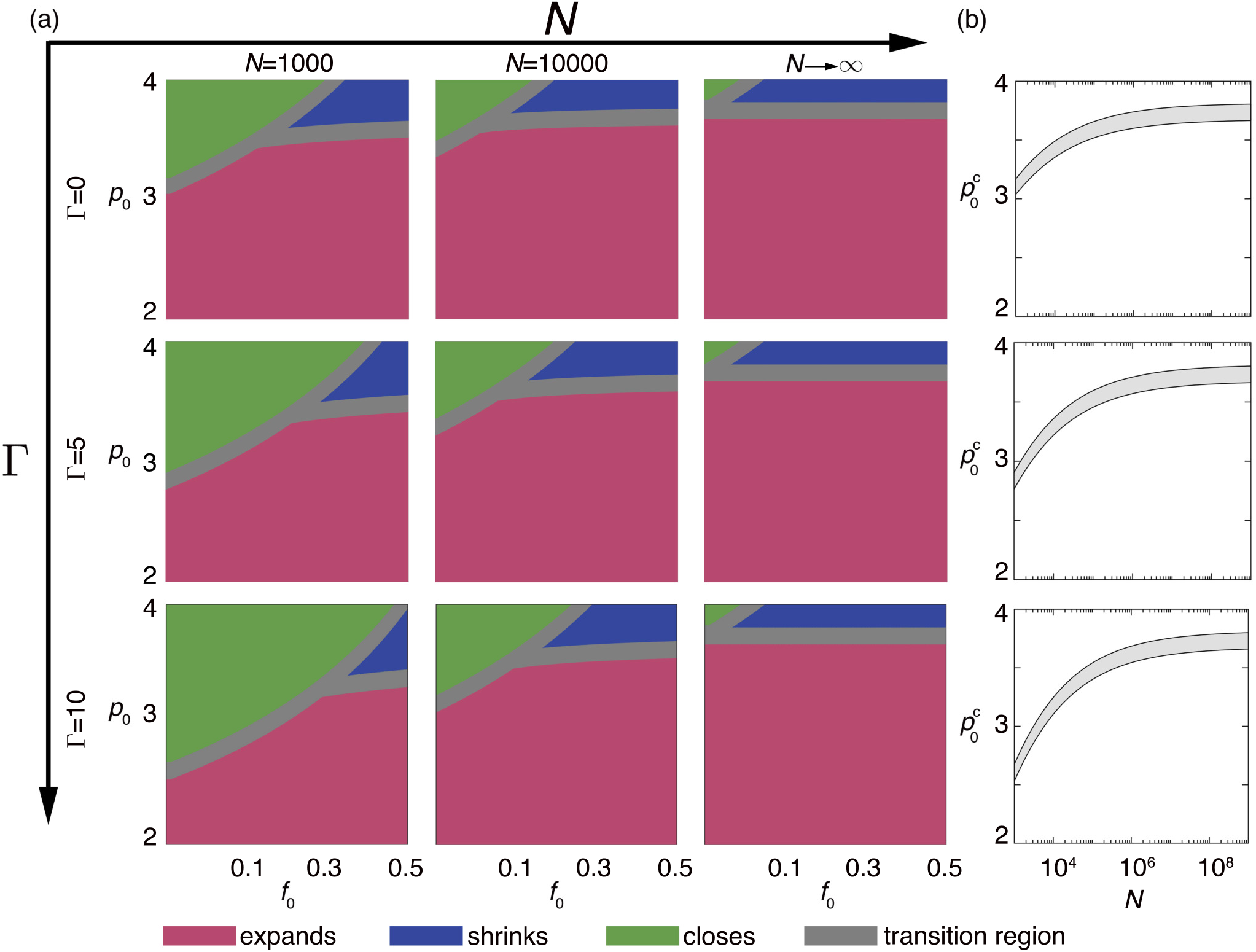
(a) Phase diagrams of wound fates as a function of preferred perimeter (*p*_0_) and initial wound fraction (*f*_0_). The three wound fates are: closure (green), shrinkage (blue), and expansion (magenta). The grey-coloured region represents the region in phase-space, where the prediction of our model is not uniform for 3.67 *< c <* 3.81. Different phase diagrams represent different values of cell number (*N* ) and adimensional tissue-wound interfacial tension (Γ). (b) Critical preferred perimeter for wound closure 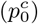 as a function of the total number of cells in the monolayer (*N* ) for the Γ = 0, 5, and 10 respectively. The upper and lower bounds of the transition region represent the value calculated for *c* = 3.81, and *c* = 3.67 respectively.

**FIG. 3.**
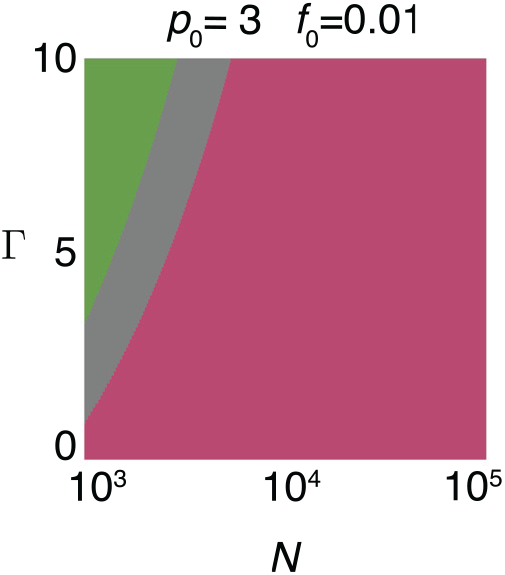
Phase diagram showing wound fates as a function of tissue-wound interfacial tension (Γ) and tissue size (*N* ) for a given value of preferred perimeter (*p*_0_) and initial wound fraction (*f*_0_). The two wound fates are: closure (green) and expansion (magenta). The grey-coloured region represents the region in phase-space, where the prediction of our model is not uniform for 3.67 *< c <* 3.81.

The phase-space is characterized by a set of features [Fig. 2(a), Fig. 3]. For a tissue with specific *N* and Γ, if *p*_0_ is lower than a critical value 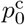 , no wound closes, and all of them rupture. For 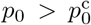 , wounds upto a certain size (*f*_0_) closes. As the *p*_0_ increases (the tissue becomes more adhesive), wounds with larger area fraction closes. For a positive Γ, the internal adhesion of the tissue gets assisted by purse-string tension. As a result,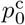 decreases, and wounds of the same size close at a lower *p*_0_. However, the effect of these mechanisms reduces as the tissue becomes larger and larger because they act only at the wound perimeter and the ratio of the wound perimeter to the area of the wound decreases (for the same wound area fraction). The critical value 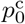 increases with system size *N* as shown in Fig. 2(b). In a tissue of a specific mechanical property, wounds of a given area fraction *f*_0_ close in a smaller size tissue, but not in a larger size tissue [Fig. 3]. In fact, all the phase diagrams collapse at the limit *N* → ∞. Irrespective of the value of Γ, wounds close only when *p*_0_ *> c* [Fig. 2(a)].

Our theoretical analysis makes some interesting predictions about the geometric limit of wound closure. Specifically, the phase diagrams shown in Fig. 2(a) suggest that for a tissue of specific size and specific mechanical properties, only wounds upto a certain area fraction close and beyond that it does not close. On the other hand, wounds of a certain area fraction close in a smaller size tissue but not in a larger size tissue [Fig. 2(a), Fig. 3].

### Experimental Validation

To validate if our theoretical prediction is truly manifested in an epithelial monolayer, we design an experimental assay. In the natural process of tissue healing, multiple mechanisms collaborate. For instance, on an adherent substratum beneath the epithelium, cells are encouraged to infiltrate through lamellipodia-driven crawling, making migration the primary mechanism responsible for gap closure [8]. Conversely, on a non-adherent substratum, cells exhibit coordinated acto-myosin purse-string driven contractile motions, closing wounds without requiring migration [53, 54]. This phenomenon is observable using micropatterned surfaces that selectively apply adhesive coatings like collagen or fibronectin around the target wound area [12]. Additionally, in the traditional scratch assay [55], which removes cells along with patches of the underlying adherent substrate, wound closure occurs through a mixed mode involving both acto-myosin purse-string contractility and cell migration. Moreover, cell proliferation significantly contributes to wound healing in living tissues, particularly over extended time scales [56].

At the tissue level, these mechanisms operate in a coordinated manner, involving numerous specific regulatory pathways that extend beyond the scope of the model presented in this study [57]. Our model primarily concentrates on the mechanical responses of the tissue, omitting explicit incorporation of cell migration or proliferation. In this context, we refer to previous experiments that used micro-patterned surfaces to identify limitations on wound closure in the absence of cell crawling [12]. In these experiments, actin-based contractility predominantly facilitates wound closure over non-adherent gaps. Moreover, this research identified dimensions of wound sizes beyond which closure was either completely or partially impeded, while below which complete recovery was observed similar to the prediction of our model. This alignment with the predictions from our theoretical model suggests analogous geometric constraints on wound closure controlled by both tissue size and wound fraction.

Motivated by these parallels between past experimental work and our present model, we design an experimental assay [Fig. 4(a)] to validate our predictions about the wound healing processes with variable cell number and variable wound size relative to the tissue. In each of the assays, we seeded homogeneous single cell suspensions of MDCK epithelial cells in a petridish coated with collagen monomers except for a region shielded by a sacrificial agar droplet of a specific size. The seed suspension rapidly proliferates generating a tightly packed confluent monolayer uniformly surrounding the agar droplet, which simulates the “wound” area. After the monolayer is formed, the agar droplet is removed and we can observe the fate of the wound over several hours using confocal microscopy. This setup eliminates the need for destructive procedures such as laser ablation and allows us to precisely control the initial wound area (by controlling the size of the agar droplet) as well as the tissue size (by controlling the size of the petridish). We note that while the lack of adherent collagen coatings over the wounded area appears to largely prohibit cellular migration, it is not possible to completely eliminate this aspect from the ensemble tissue. Further, to minimize the contributions from proliferative additions to the tissue, we restrict our observations within a 24-hr window, which is on the order of typical cell division times for the MDCK.

**FIG. 4.**
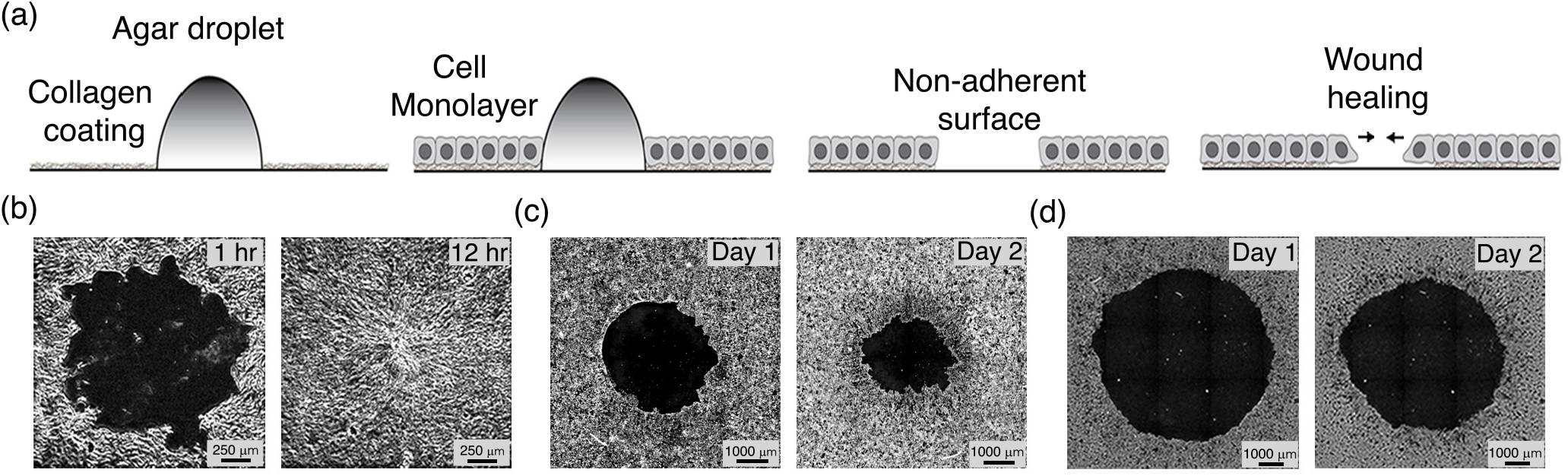
(a) Schematic of the experimental setup to mimic a wound in a confluent epithelial monolayer. Panels (b)-(d) show micrographs of initial (left) and final (right) state of the wound corresponding to three different sets of geometric parameters. For the micrograph shown in (b)-(d) the initial area fractions (*f*_0_) are 0.009, 0.009, and 0.145, respectively. The diameters of the petri dishes (proportional to 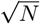 ) in panels (b)-(d) are 12 mm, 35 mm, and 12 mm, respectively.

In Fig. 4(b)-(d), we show three examples of our assay. By adjusting the size of agar droplet within a given petri dish, we have the ability to alter the wound fraction (*f*_0_), whereas altering the size of the dish itself affects the tissue size (*N* ). We observe that our experimental assays closely match predictions from our theory. For the same tissue size *N* , only wounds with a smaller *f*_0_ close, whereas for the same *f*_0_, wounds close only in a tissue with smaller *N* . After observing this qualitative agreement between the theoretical prediction and experimental observation, we investigate for what range of mechanical parameters our theoretical model would make the same quantitative prediction. Specifically, we ask for what values of Γ and *p*_0_ in a tissue of size *N* = 1.13 × 10^6^, a wound with *f*_0_ = 0.009 closes but a wound with *f*_0_ = 0.145 doesn’t close. We identify and show the range in Fig. 5. Additionally, we extend this inquiry beyond our own experiments presented in this paper and examine a similar experimental outcome reported in [12]. Specifically, we investigate a tissue of size *N* = 6400, where a wound with *f*_0_ = 0.049 closes but a wound with *f*_0_ = 0.110 does not close. The corresponding parameter region is also illustrated in Fig. 5. We estimate cell numbers by measuring the characteristic cell area in a confluent epithelial monolayer, which is approximately 100 *μ*m^2^. Interestingly, the mechanical parameter space exhibits a narrow intersection, where our model accurately predicts the experimental outcomes from both studies. Furthermore, we would like to highlight that the values estimated in several studies for Γ and *p*_0_ on living tissues actually fall in that narrow intersection, which is specifically marked by the black rectangle [37, 47, 52, 58–62]. Altogether, these results suggest that our model robustly predicts the mechanical response of a living tissue to formation of a wound when the effect of cell proliferation and cell crawling is minimal.

**FIG. 5.**
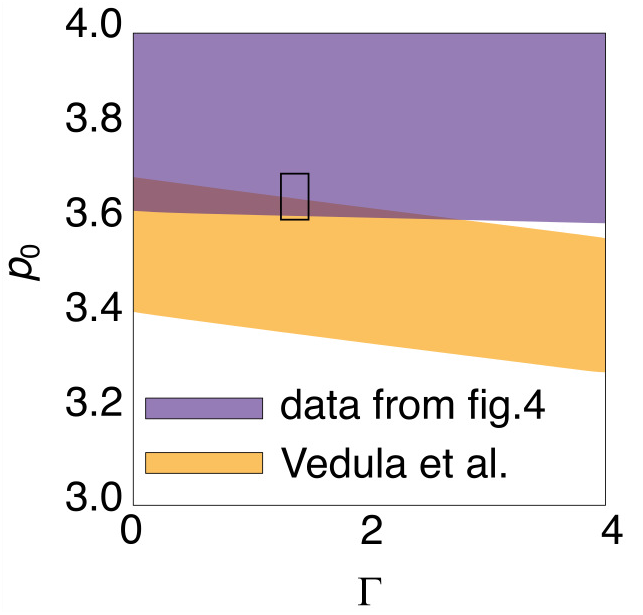
The coloured regions show the range of mechanical parameters (*p*_0_ and Γ) for which the experimental observation quantitatively matches with the theoretical prediction. i.e., for a given tissue size (*N* ), a smaller wound of a specific size (*f*_0_) exhibits closure, whereas a larger wound of another specific size fails to close. The region marked in purple and yellow shows the parameter range for the experiments reported in our work and the work by Vedula et. al respectively [12]. The black rectangle represents the range of *p*_0_ and Γ values reported by previous studies on living tissues [37, 47, 52, 58– 62].

## DISCUSSIONS

In this article, we introduce a mathematical framework to predict whether a wound will close, shrink, or rupture in an epithelial monolayer. The framework relies on a coarse-grained mathematical representation of the vertex model description of the tissue and does not need any computationally intensive dynamic simulations. Our framework studies the effects of geometrical properties of the wound and mechanical properties of the tissue in governing wound closure. The geometrical properties include tissue size and initial area fraction of the wound, while acto-myosin cortex contractility, cell-cell adhesion, and purse-string action constitute the mechanical properties of the tissue. Specifically, we quantitatively establish that within a tissue, cell-cell adhesion must meet a certain critical value 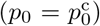 for wound closure to occur. Beyond that *p*_0_, we identify that wounds with an initial area fraction (*f*_0_) not exceeding a finite value close. Additionally, we observe that wound closure is influenced positively by tissue-mediated mechanisms like the purse-string action (Γ), but hindered by larger tissue sizes (*N* ).

Furthermore, we validate several of our model’s predictions by designing an experimental setup that allows precise control of wound-related geometric properties, including the initial area fraction of the wound (*f*_0_) and tissue size (*N* ). We confirm that within tissues of the same size, wounds with a higher initial area fraction fail to close. Also, wounds with the same area fraction do not close in larger tissues. Furthermore, we quantitatively show that for the mechanical parameters reported for living tissues, our theoretical model robustly predicts our experimental observation and a similar study reported in [12]. However, to further assess our predictions about the effect of mechanical properties such as cell-cell adhesion, and tissue-initiated mechanisms like purse-string action, future experiments are needed that will require controlled perturbation of these properties by genetic or biochemical approaches.

Through the integration of theoretical modeling and experimental validation, this work reveals that the geometrical and mechanical properties of an epithelial tissue collectively decide the fate of a wound. In future, this model can be adapted to take into account events that dynamically change the number of cells in the tissue, such as tissue proliferation through cell division or tissue pruning via cell death. On the other hand, one can use this study as a basis to explore how a non-living viscoelastic material responds to the formation of a rupture and compare it with the response of a healing tissue to the formation of a wound. Recent works that relate tissue rheological properties to vertex model parameters are especially relevant in this context [33, 63–65]. Altogether, this work serves as a foundational step towards a deeper understanding of the intricate interplay and synergy among diverse mechanical and geometrical properties of tissues in the process of wound healing.

## AUTHOR CONTRIBUTIONS

TB and SD designed the project. NKB, SD and TB did the theoretical formulation and analysis. MS performed the experiments. TB guided the overall project. All authors contributed in writing the manuscript.

## ACKNOWLEDGEMENTS

NKB and MS acknowledge TIFR Graduate student program. TB acknowledges SERB CRG/2021/008869/EXP grant for research funding. TB also acknowledges NCBS-TIFR for intramural funding.

